# Enzymatic modulation of the pulmonary glycocalyx alters susceptibility to *Streptococcus pneumoniae*

**DOI:** 10.1101/2024.01.03.573996

**Authors:** Cengiz Goekeri, Kerstin A.K. Linke, Karen Hoffmann, Elena Lopez-Rodriguez, Vladimir Gluhovic, Anne Voß, Sandra Kunder, Andreas Zappe, Sara Timm, Alina Nettesheim, Sebastian M.K. Schickinger, Christian M. Zobel, Kevin Pagel, Achim D. Gruber, Matthias Ochs, Martin Witzenrath, Geraldine Nouailles

**Author notes:** Equal contribution (Cengiz Goekeri, Kerstin A.K. Linke). **Corresponding author:** Geraldine Nouailles, Dr.-Ing., Charité - Universitätsmedizin Berlin, Corporate Member of Freie Universität Berlin and Humboldt-Universität zu Berlin, Department of Infectious Diseases, Respiratory Medicine and Critical Care Charitéplatz 1, 10117 Berlin, Germany; /. **Author Contributions:** CG, KAKL, KH, GN performed experiments, analyzed and interpreted the data. AN and SMKS performed experiments. ELR, VG, ST performed EM analysis, and interpreted the data. MO interpreted and supervised EM studies. AV, SK performed histopathology and interpreted the data. ADG interpreted and supervised histopathology. AZ performed HPLC analysis and interpreted the data. KP interpreted and supervised HPLC analysis. CMZ provided resources, and interpreted Chip data. MW interpreted data. GN, CG and KAKL drafted the manuscript. KH, ELR, AV, SK, AZ and ADG wrote sections of the manuscript. GN conceived the study and supervised the work. All authors revised and edited the manuscript and approved the final submitted version for publication.

## Abstract

The pulmonary epithelial glycocalyx is rich in glycosaminoglycans such as hyaluronan and heparan sulfate. Despite their presence, the precise role of these glycosaminoglycans in bacterial lung infections remains elusive. To address this, we intranasally inoculated mice with *Streptococcus pneumoniae* in the presence or absence of enzymes targeting pulmonary hyaluronan and heparan sulfate, followed by characterization of subsequent disease pathology, pulmonary inflammation, and lung barrier dysfunction. Enzymatic degradation of hyaluronan and heparan sulfate exacerbated pneumonia in mice, as evidenced by increased disease scores and alveolar neutrophil recruitment. However, targeting epithelial hyaluronan further exacerbated systemic disease, indicated by elevated splenic bacterial load and plasma levels of pro-inflammatory cytokines. In contrast, enzymatic cleavage of heparan sulfate resulted in increased bronchoalveolar bacterial burden, lung damage and pulmonary inflammation in mice infected with *Streptococcus pneumoniae*. Accordingly, heparinase-treated mice also exhibited disrupted lung barrier integrity as evidenced by higher alveolar edema scores and vascular protein leakage into the airways. This finding was corroborated in a human alveolus-on-a-chip platform, confirming that heparinase treatment also disrupts the human lung barrier. Notably, enzymatic pre-treatment with either hyaluronidase or heparinase also rendered human epithelial cells more sensitive to pneumococcal-induced barrier disruption, as determined by transepithelial electrical resistance measurements, consistent with our findings in murine pneumonia. Taken together, these findings demonstrate the importance of intact hyaluronan and heparan sulfate in controlling pneumococcal virulence, pulmonary inflammation, and epithelial barrier function.

## INTRODUCTION

Community-acquired pneumonia (CAP) is associated with high morbidity and mortality despite vaccination and antibiotic use, with *Streptococcus pneumoniae* detected as the most frequent bacterial pathogen in CAP patients (1–3). Disease progression may result in severe CAP, defined by acute respiratory failure and the need for mechanical ventilation (4–6). Acute lung injury is characterized by hypoxemia, enhanced pulmonary vascular permeability and inflammation-mediated lung injury (7, 8).

One factor involved in acute lung injury that remains incompletely understood is the functional role of the glycocalyx in the context of pulmonary inflammation (9). The glycocalyx is a functional layer of glycoproteins and proteoglycans anchored to the plasma membrane of virtually all cells (10), including lung epithelial cells (11). The proteoglycans are bound to the cell membrane via core proteins containing a transmembrane domain and have long unbranched glycosaminoglycan (GAG) side chains consisting of polysaccharides with repeating disaccharide units. The most common GAGs in the lung glycocalyx are anticipated to be chondroitin sulfate, heparan sulfate and hyaluronan (12).

Unlike the glycocalyx of the lung epithelium, the role of the endothelial glycocalyx in pulmonary capillaries has been widely studied: It serves as a mechanosensor and signal transducer (13–15), regulates vascular permeability (16, 17) and binds anticoagulant mediators (18), growth factors and chemokines (19, 20). While these functions are of elementary importance, many of them are closely linked to vascular blood flow, leaving the functional role of the alveolar epithelial glycocalyx unclear. It has been postulated that it may be involved in maintaining a functioning air-blood barrier and that shed heparan sulfate may bind inhaled pathogens as a decoy receptor (9). A limited number of studies have investigated its role during lung inflammation, demonstrating alveolar heparan sulfate shedding in mice following LPS-induced lung injury, as well as in acute respiratory distress syndrome (ARDS) patients. In both cases, GAG shedding was correlated with increased lung permeability (9, 21).

The role of the epithelial glycocalyx during bacterial pneumonia is of particular interest, as pathogenic bacteria produce a variety of virulence factors to modify host carbohydrates and improve adherence (22, 23). In particular, *S. pneumoniae* produces the virulence factor hyaluronidase (24, 25) and may thus be able to cleave one of the main GAGs of the glycocalyx. In addition, a recent study demonstrated that the pneumococcal exotoxin pneumolysin contributes to shedding of the glycocalyx from alveolar epithelial cells (26).

Collectively, however, the involvement of microbial and host-mediated GAG processing in the orchestration of pulmonary inflammation and disease pathology remains poorly understood. Here, we utilized intranasal delivery of heparinase and hyaluronidase to degrade two major glycocalyx GAGs during *S. pneumoniae* infection in a murine model of pneumococcal pneumonia.

## MATERIALS AND METHODS

### Mice and Housing

Specific pathogen-free female C57BL/6J mice were obtained from Janvier Labs (Le Genest-Saint-Isle, France) at 8-10 weeks of age and housed under pathogen-free conditions for at least 5 days before experimental treatments. Mice were kept in groups of 3 to 4 in individually ventilated cages with a 12 h day-night cycle and had free access to food and water. All experiments were performed in compliance with national and international guidelines for the care and humane use of animals and approved by the relevant state authority, Landesamt für Gesundheit und Soziales (LAGeSo) Berlin, Germany (No. G0099/21).

### Infection, monitoring and dissection

Animals were anesthetized by intraperitoneal injection of ketamine-xylazine (93.75 mg/kg ketamine, Bela-Pharm, Vechta, Germany and 15 mg/kg xylazine, CP-Pharma, Burgdorf, Germany) and infected transnasally in an upright position with a final volume of 20 µl 1×PBS containing 5 × 10^6^ CFUs of *S. pneumoniae* serotype 3 (ST3) with or without addition of either Hyaluronidase from bovine testes Type I-S (H3506 Sigma-Aldrich, St. Louis, US; 90 units (unit definition: One unit will degrade 0.75 μg of the polysaccharide hyaluronic acid per minute at pH 5.35 at 37 °C (as measured by turbidimetric absorbance (λ: 600 nm) when complexed with BSA after 45 minutes)) dissolved in 1×PBS) or Heparinase I and III Blend from *Flavobacterium heparinum* (H3917 Sigma-Aldrich, St. Louis, US; 15 Sigma units (unit definition: One Sigma unit will form 0.1 µmol of unsaturated uronic acid per hour at pH 7.5 at 25 °C) dissolved in 1×PBS). Control (sham-infected) mice received 20 μl PBS with or without the addition of enzymes. Corneas were protected under anesthesia by Thilo-Tears Gel (Alcon Deutschland GmbH, Aschaffenburg, Germany). Clinical parameters were recorded and behavioral signs of disease scored (Table E1) prior to the start of the experiment and every 12 h from 24 h post-infection (hpi). Dissection of the animals was performed at 2 hpi for electron microscopic analysis and 48 hpi for all other analyses, under terminal ketamine-xylazine anesthesia (i.p. injection; 200 mg/kg of body weight ketamine and 20 mg/kg of body weight xylazine) and exsanguination after loss of the intertoe reflex.

### Data analysis

Results were analyzed using GraphPad Prism 9 (GraphPad Software, Boston, USA). Statistical significance was determined as described in the figure legends. All graphs depict mean ± SEM unless otherwise stated; p < 0.05 was considered statistically significant.

All other details are described in the extended material and methods section of the online supplement.

## RESULTS

### Enzymatic targeting of epithelial glycosaminoglycans aggravates pneumococcal pneumonia

To improve our understanding of epithelial glycocalyx function during bacterial pneumonia, we targeted common glycocalyx GAGs enzymatically with heparinase or hyaluronidase in a murine pneumococcal pneumonia model, by adding the enzymes directly to the bacterial inoculum. C57BL/6J mice were transnasally infected with a mixture of *S. pneumoniae* ST3 and either PBS only (infected-only control) or PBS containing either heparinase (15 Sigma units), or hyaluronidase (90 units). Mock-infected controls received PBS with or without enzymes but without the addition of bacteria. Clinical status was monitored at 24, 36 and 48 hpi and animals were sacrificed for further analysis at 48 hpi, when pneumonia had developed. In addition, 3 animals in each group were sacrificed at 2 hpi for electron microscopic imaging (Fig. 1A). Clinical monitoring for signs of pneumonia included measurements of body temperature (Fig. 1B), weight (Fig. 1C), survival (Fig. 1D) and behavioural disease score (Fig. E1A, Table E1). Unlike mock-infected animals, mice that were infected with *S. pneumoniae* displayed loss of body temperature and weight over infection time (Fig. 1B, C). Addition of hyaluronidase or heparinase to the infection inoculum aggravated the disease course: Body temperature loss was significantly more pronounced by 36 hpi, whilst body weight loss remained similar over time in infected mice, independent of treatment (Fig. 1B, C). At 48 hpi, the disease score of enzyme-treated mice worsened significantly towards inactivity, poorer grooming and labored breathing (Fig. E1A). Accordingly, 6 of 19 and 2 of 16 infected mice from the heparinase and hyaluronidase group, respectively, had to be euthanized at humane endpoints set by the regulatory authority prior to 48 hpi, compared to only 1 of 15 mice from the infected group that received no enzymes (Fig. 1D).

**Figure 1:**
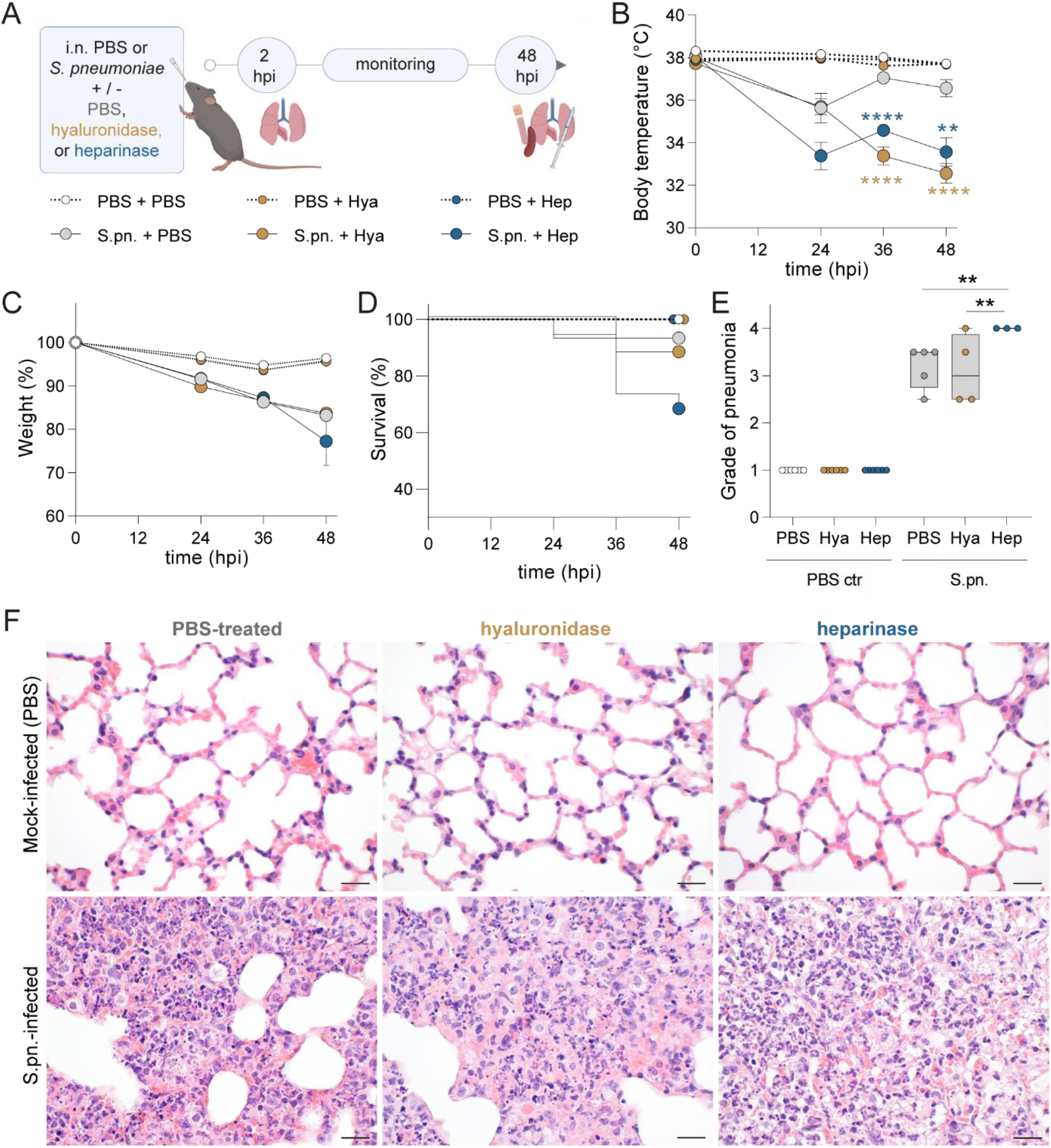
Enzymatically targeting pulmonary glycosaminoglycans worsens symptoms and severity of pneumococcal pneumonia. Mice were infected i.n. with *S. pneumoniae* (S.pn.) or PBS (PBS ctr) with addition of hyaluronidase (Hya) or heparinase (Hep) or PBS as control and sacrificed at 48 hpi. (**A**) Experimental layout. Graphs display (**B**) body temperature (°C), (**C**) weight change (%) and (**D**) survival of experimental animals over the infection course. (**B**, **C**) Mixed-effect analysis, Dunnett’s multiple comparisons test, tested against S.pn. + PBS, test results displayed for S.pn. groups; n = 15-19 mice per group. (**D**) Log rank (Mantel-Cox) test. (**E**) Histopathological degree of pneumonia ranging from (0) absent to (4) severe at 48 hpi. n = 3 – 6 mice per group. Two-way ANOVA, Tukey’s multiple comparison test. Data are displayed as box plots, middle line displays median, box indicates first and third quartile, and whiskers minimum to maximum. Significant differences are only shown between infected control vs. treated groups (** p ≤ 0.01 and **** p ≤ 0.0001). (**F**) Hematoxylin and eosin staining of murine lungs. While mock infection resulted in almost no lesions across all groups (upper panel), all infected groups developed diffuse alveolar damage with highest severity, density, and damage in heparinase-treated mice (H&E, original magnification 600x. Bars equal 20 µm).

Histopathology scores were discriminative between infected and mock-infected mice. Among the enzymatically-treated groups, heparinase-treated mice had the most severe pneumonia scores (Fig. 1E, Table E2). All mock-infected, including enzyme-only treated groups, developed almost no histopathological changes (Fig. 1F, upper panel). In contrast, infection resulted in mostly moderate, multifocal, purulent bronchoalveolar pneumonia (Fig. 1F, lower panel) with diffuse alveolar damage.

Next, we assessed the bacterial burden and dissemination in *S. pneumoniae*-infected mice. Enzyme-treated groups displayed higher bacterial loads in BAL and lungs compared to the infected PBS-treated group (Fig. 2A, B). Dissemination to the vasculature was mildly elevated in all groups, but only hyaluronidase-treated mice displayed significantly higher bacterial burden in the spleen compared to PBS-treated infected mice (Fig. 2C, D). To probe whether enzyme treatment directly affected bacterial growth, we grew *S. pneumoniae* ST3 in liquid culture in the presence of heparinase (30 Sigma units/ml) or hyaluronidase (180 units/ml) and measured bacterial growth at OD_600nm_ until stationary phase. We observed no difference in growth compared to the control group cultured without enzymes (Fig. E1B). In addition, we spread inoculum samples (750 Sigma units/ml heparinase, 4500 units/ml hyaluronidase) on blood agar plates and counted the colony-forming units (CFUs) after overnight incubation. Again, we observed no difference between samples containing enzymes and controls (Fig. E1C).

**Figure 2:**
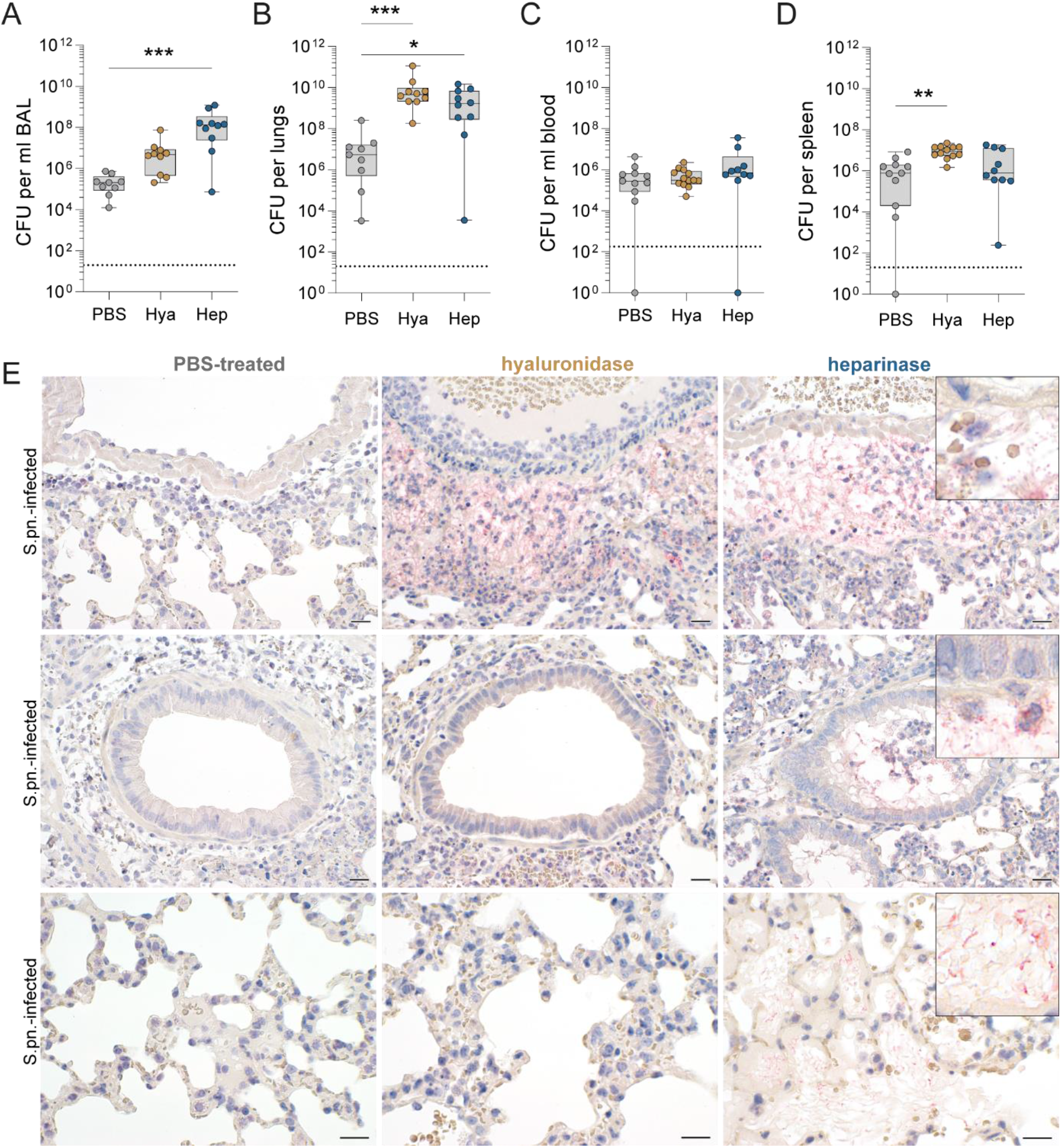
Enzymatic targeting of epithelial glycocalyx alters susceptibility to streptococcal pneumonia. Mice were infected i.n. with *S. pneumoniae* (S.pn.) with addition of hyaluronidase (Hya) or heparinase (Hep) or PBS as control and sacrificed at 48 hpi. (**A – D**) Colony-forming units (CFU) for (**A**) BAL, (**B**) lungs, (**C**) blood and (**D**) spleen. Kruskal-Wallis and Dunn‘s multiple comparison test; n = 9 – 13. Data are displayed as box plots, middle line displays median, box indicates first and third quartile, and whiskers minimum to maximum. Dotted lines indicate limit of detection (LOD), BAL: 20 CFU/ml; lungs and spleen: 20 CFU/organ, blood: 200 CFU/ml. Samples below LOD were set to 1 or 1/ml. * p ≤ 0.05, ** p ≤ 0.01, and *** p ≤ 0.001 (**E**) Immunohistochemical visualization of *S. pneumoniae* (New Fuchsin, red) in lungs from S.pn-infected and PBS-treated (left panel), hyaluronidase-treated (middle panel) or heparinase-treated (right panel) mice with hematoxylin (blue) as counterstain. Top and middle panel, original magnification 400x. Bars equal 20 µm. Bottom panel, original magnification 600x. Bars equal 20 µm. Original magnification of inserts: 1000x.

As streptococcal burden in organs pointed towards biased localization between groups, we visualized pulmonary *S. pneumoniae* by New Fuchsin staining |(red) and immunohistochemistry (Fig 2E). At 48 hpi, PBS-treated mice had no free bacteria in perivascular spaces that were infiltrated with neutrophils (top), in or around airways (middle), or in the respiratory parenchyma (bottom). In contrast, both hyaluronidase- and heparinase-treated mice had abundant free streptococci in the perivascular and peribronchial spaces. In addition, heparinase-treated mice also had numerous intrabronchiolar and intraalveolar free streptococci (right, middle and bottom panel) (Fig. 2E and Fig. E1D).

### Neutrophil recruitment and inflammation are enhanced following enzymatic targeting of epithelial glycosaminoglycans

Immunohistopathology revealed streptococci targeted by professional phagocytes (Fig. 2E). We thus quantified the presence of innate immune cells at 48 hpi, using flow cytometric analysis of BAL cells (gating strategy in Fig. E2). Increased numbers of CD45^+^ leukocytes, neutrophils and Ly6C^hi^ monocyte-derived inflammatory macrophages were detected in all infected groups. Addition of hyaluronidase and heparinase increased neutrophil numbers further than infection alone. In contrast, recruitment of Ly6Chi iMs remained at PBS treatment level for heparinase-treated mice and was significantly lower in hyaluronidase-treated mice (Fig. 3A – C). Upon infection, neutrophils represented over 70% and Ly6C^hi^ inflammatory macrophages around 2 – 4% of the alveolar leukocytes; the proportion of neutrophils was also significantly enhanced in mock-infected heparinase-treated mice (Fig. 3D – G). We did not detect any significant differences in the numbers of alveolar macrophages in response to enzyme treatment at this stage of infection, although the overall proportion of alveolar macrophages was reduced in all infected mice, and significantly so in heparinase treated mice, likely as a result of the massive neutrophil influx (Fig. E3A – C). Notably, alveolar concentrations of interleukin (IL-)17A, which is associated with epithelial activation towards neutrophil recruitment, as well as of the neutrophil-chemoattractants CXCL1 and 5, were significantly increased upon heparinase treatment in infected mice (Fig. 3H – J).

**Figure 3.**
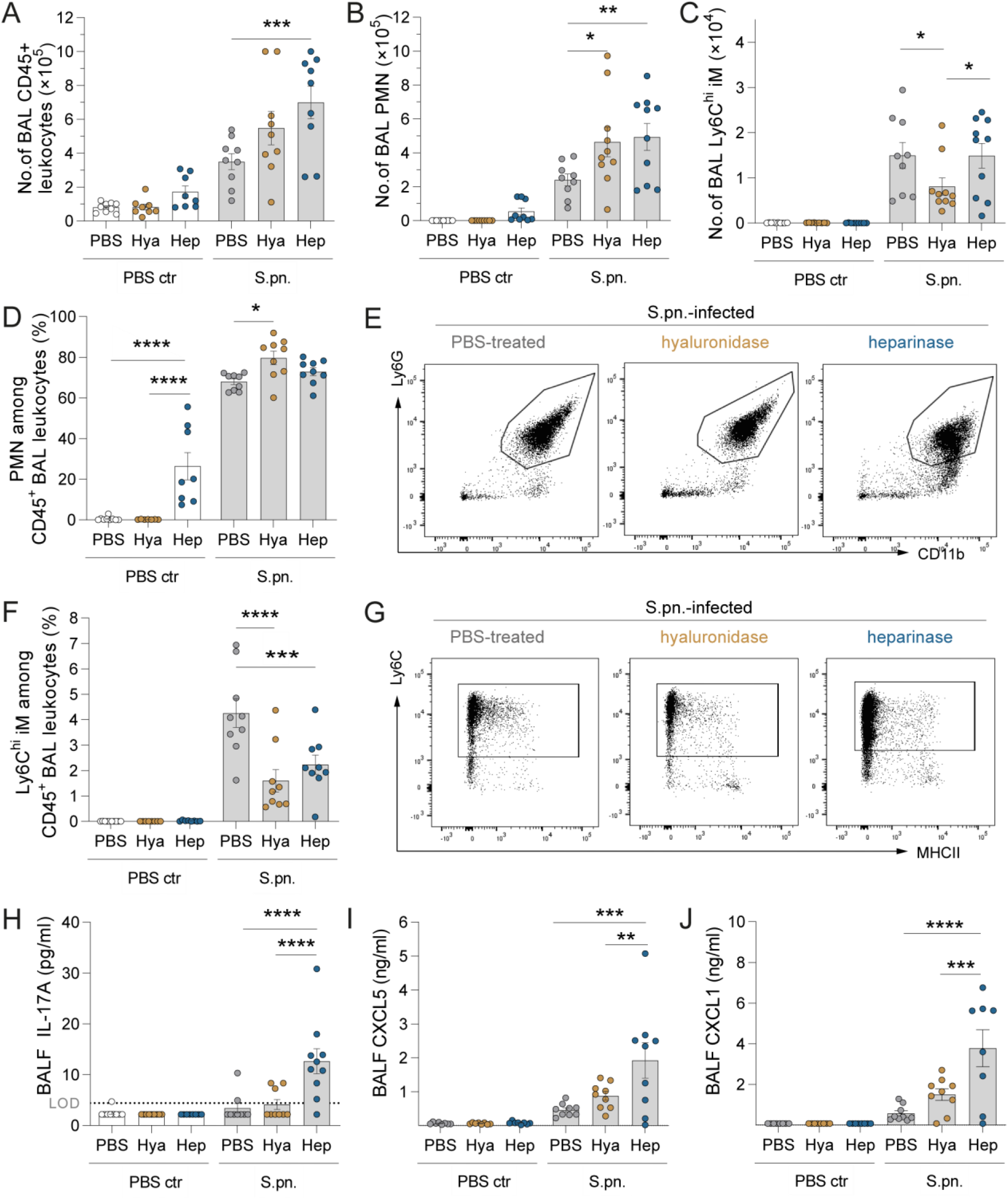
Inflammatory cell recruitment is enhanced following targeting of epithelial glycocalyx in pneumonia. Mice were infected i.n. with *S. pneumoniae* (S.pn.) or PBS (PBS ctr) with addition of hyaluronidase (Hya) or heparinase (Hep) or PBS as control and sacrificed at 48 hpi. **(A – G)** Flow cytometry-based analysis of immune cells. Total numbers of (**A**) CD45^+^ leukocytes, (**B**) polymorphonuclear neutrophils (PMN) and (**C**) Ly6C^hi^ inflammatory monocyte-derived macrophages (Ly6C^hi^ iM) in BAL were determined by use of counting beads while cell frequencies **(D – G)** were determined as percentage of CD45^+^. (**E**, **G**) Dot plots representing cellular frequencies amongst CD45^+^ leukocytes. **(H – J)** Protein concentrations of IL-17A (limit of detection (LOD) 4.42 pg/ml), CXCL5 and CXCL1 in bronchoalveolar lavage fluid (BALF) as quantified by ELISA. Dotted lines indicate limit of detection (LOD). Values below LOD were set to half LOD for statistical analysis. Two-way ANOVA and Tukey’s multiple comparisons test; n = 8 – 10. * p ≤ 0.05, ** p ≤ 0.01, *** p ≤ 0.001 and **** p ≤ 0.0001.

To gain more comprehensive insight into effects on inflammatory mediators known to play a role in pneumococcal infection, we quantified the concentrations of IL-6, IL-1β, tumor necrosis factor alpha (TNF-α), interferon gamma (IFN-γ), and chemokine ligand 2 (CCL2). Without infection, we observed no increase in BALF concentrations of any of these factors in response to enzyme treatment. Heparinase treatment of *S. pneumoniae*-infected animals resulted in significantly elevated BALF concentrations of the examined inflammatory cytokines compared to infected-only or hyaluronidase-treated animals (Fig. 4A – E). In line with this, histopathological evaluation found larger areas of inflammation in infected heparinase-treated animals (Fig. 1F and Fig. 4F). Treatment with hyaluronidase caused no or only non-significant increases in cytokine levels over infection alone, except for BALF IFN-γ (Fig. 4D). Cytokine levels in plasma were only mildly elevated upon infection for IL-6, IFN-γ, TNF-α and CCL2. Enzymatic treatment further elevated the concentrations, with significant increases found for plasma levels of IL-6, TNF-a, IFN-γ and CCL2 upon hyaluronidase treatment (Fig. E4A – E).

**Figure 4.**
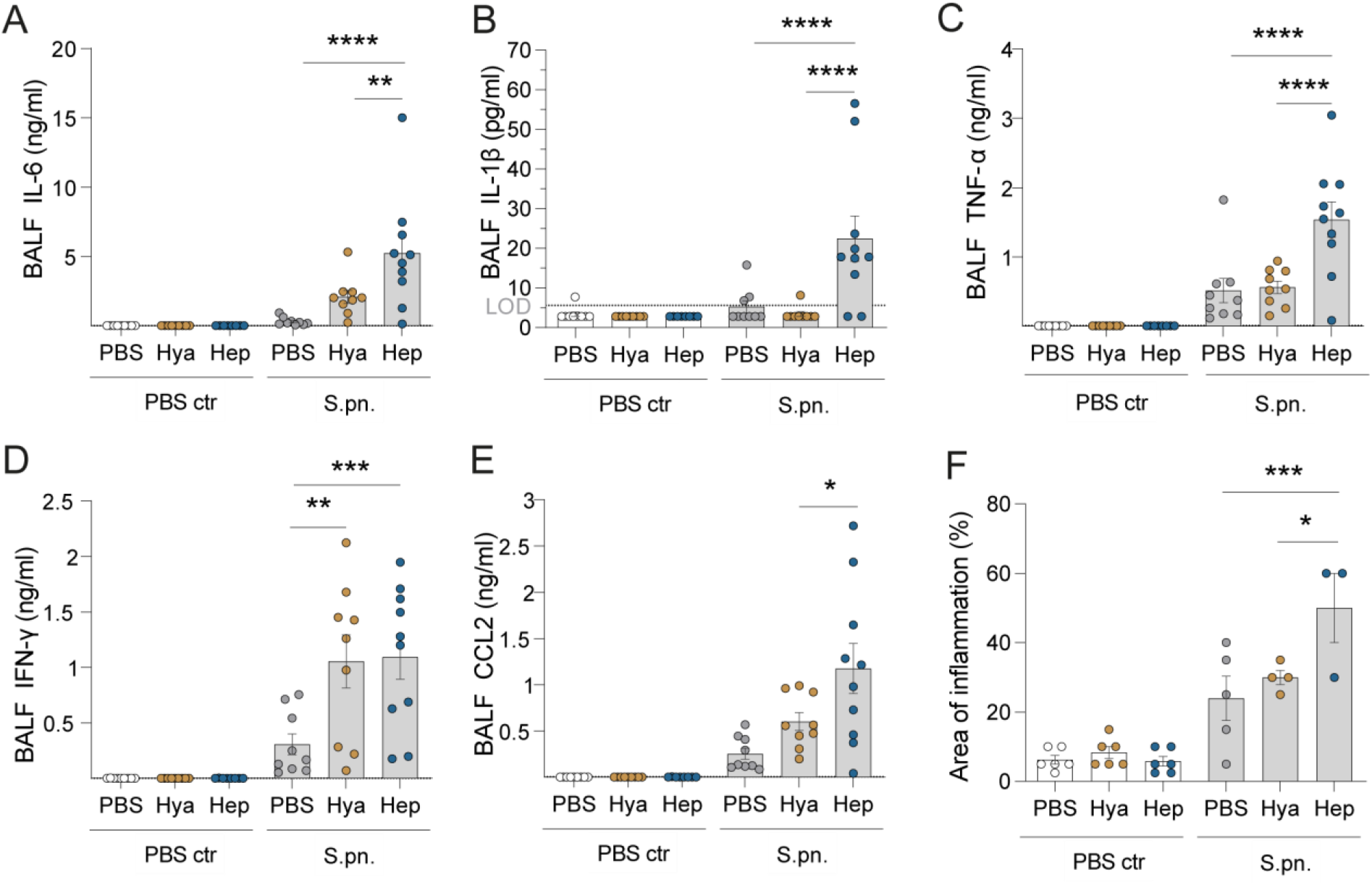
Targeting of epithelial glycocalyx enhances release of pro-inflammatory mediators. Mice were infected i.n. with *S. pneumoniae* (S.pn.) or PBS (PBS ctr) with addition of hyaluronidase (Hya) or heparinase (Hep) or PBS as control and sacrificed at 48 hpi. The concentrations of pro-inflammatory cytokines and chemokine CCL2 in bronchoalveolar lavage fluid (BALF) were measured by multiplex ELISA: (**A**) IL-6 (limit of detection (LOD) 1.72 pg/mL), (**B**) IL-1β (LOD 5.66 pg/mL), (**C**) TNF-α (LOD 13.49 pg/mL), (**D**) IFN-γ (LOD 1.06 pg/mL), (**E**) CCL2 (LOD 5.54 pg/mL). Dotted lines indicate limit of detection (LOD). Values below LOD were set to half LOD for statistical analysis. (**F**) Area of inflammation detected during histopathological examination of H&E stained lung tissue. (**A – F**) Two-way ANOVA, Tukey’s multiple comparisons test, control group vs. treated groups. (**A** – **E**) n = 8 – 10. (**F**) n = 3 – 6. * p ≤ 0.05, ** p ≤ 0.01, *** p ≤ 0.001 and **** p ≤ 0.0001.

### Streptococcal pneumonia alters the sulfation pattern of shed heparan sulfate

To test how enzymatic targeting of heparan sulfate or hyaluronan GAGs of the glycocalyx affected subsequent GAG shedding, we used the 1,9-dimethylmethylene blue (DMMB) assay to determine the concentration of sulfated GAGs in BALF at 48 hpi. Enzymatic treatment or infection alone did not elevate levels above the low concentrations observed in PBS-only controls, while heparinase treatment in combination with infection led to a significant increase in the GAG concentration present in BALF (Fig. 5A).

**Figure 5.**
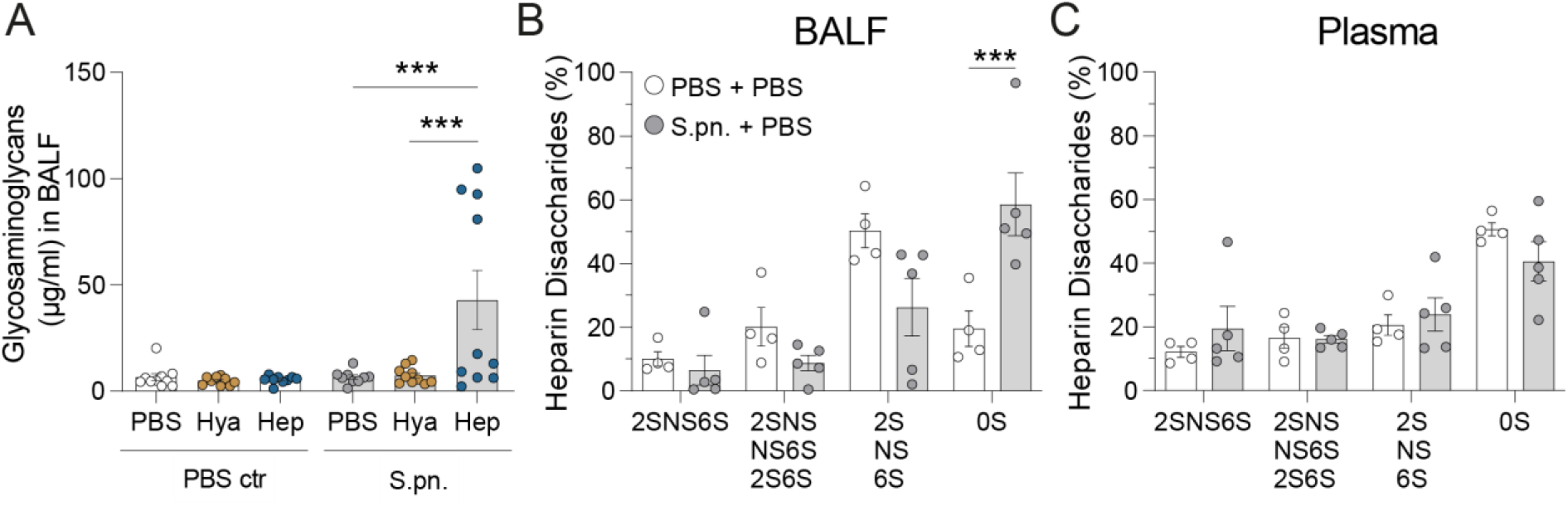
Heparin disaccharides are shed and remodeled during pneumococcal infection and following heparinase treatment in mice. Mice were infected i.n. with *S. pneumoniae* (S.pn.) or PBS (PBS ctr) with addition of hyaluronidase (Hya) or heparinase (Hep) or PBS as control and sacrificed at 48 hpi. (**A**) Glycosaminoglycan (GAG) shedding into BALF was determined by dimethylmethylene blue (DMMB) assay. Two-way ANOVA, Tukey’s multiple comparisons test; n = 9 – 10. HPLC was performed on pooled (**B**) BALF and (**C**) plasma samples to analyze sulfation of heparin disaccharides. Two-way ANOVA, Tukey’s multiple comparisons test; n = 4 – 5 (pooled from 2 – 3 independent samples). HexA-GlcNAc = 0S, HexA(2S)-GlcNAc = 2S, HexA-GlcNS = NS, HexA-GlcNAc(6S) = 6S, HexA(2S)-GlcNAc(6S) = 2S6S, HexA(2S)-GlcNS = 2SNS, HexA-GlcNS(6S) = NS6S, HexA(2S)-GlcNS(6S) = 2SNS6S. *** p ≤ 0.001.

In order to detect changes in the sulfation status of shed heparan sulfate, we determined the proportions of different disaccharides using high-performance liquid chromatography (HPLC) of full-length heparan sulfate derived from BALF (Fig. 5B) and plasma (Fig. 5C). Infection led to a marked increase in unsulfated (0S) heparin disaccharides detected in BALF (Fig. 5B), while mono-(2S, NS, 6S), di-(2SNS, NS6S, 2S6S) or tri-sulfated (2SNS6S) heparin disaccharide fractions remained similar between infected and sham groups. By contrast, in plasma, the proportions of heparin disaccharide fractions did not change significantly between infected and sham groups (Fig. 5C). It should be noted that small GAG fragments already present in the BALF, such as those generated by hyaluronidase and heparinase activity, are currently not detectable because of method limitations. For these reasons, we restricted the analysis to groups without prior enzyme treatment.

### Disruption of epithelial glycocalyx aggravates infection-triggered damage of the lung barrier

Next, we examined how enzymatic glycocalyx modulation affected lung barrier permeability to macromolecules, as severe pneumonia is characterized by breakdown of the air-blood barrier. At 48 hpi, BALF from infected animals contained elevated levels of protein, which were significantly higher in the heparinase group. Indeed, even in the non-infected animals, heparinase caused a small, insignificant increase in protein levels compared to PBS only (Fig. 6A). As expected, perivascular edema was minimal in all non-infected groups but moderate to high in the infected groups (Fig. 6B). In addition, the infected groups also showed low-to moderate-grade alveolar edema (Fig. 6C). Fluid accumulation in the adventitia was accompanied by extravasation of neutrophils and erythrocytic leakage into the perivascular space (Fig. 6D, upper panel). Notably, alveolar edema and alveolar haemorrhage was most pronounced in the heparinase-treated mice (Fig. 6D, lower panel). Taken together, combination of *S. pneumoniae* infection with heparinase treatment led to the most severe edema, inflammation and tissue damage when compared to the other groups. In addition, only heparinase treatment resulted in intra-alveolar haemorrhage (Fig. 6D).

**Figure 6.**
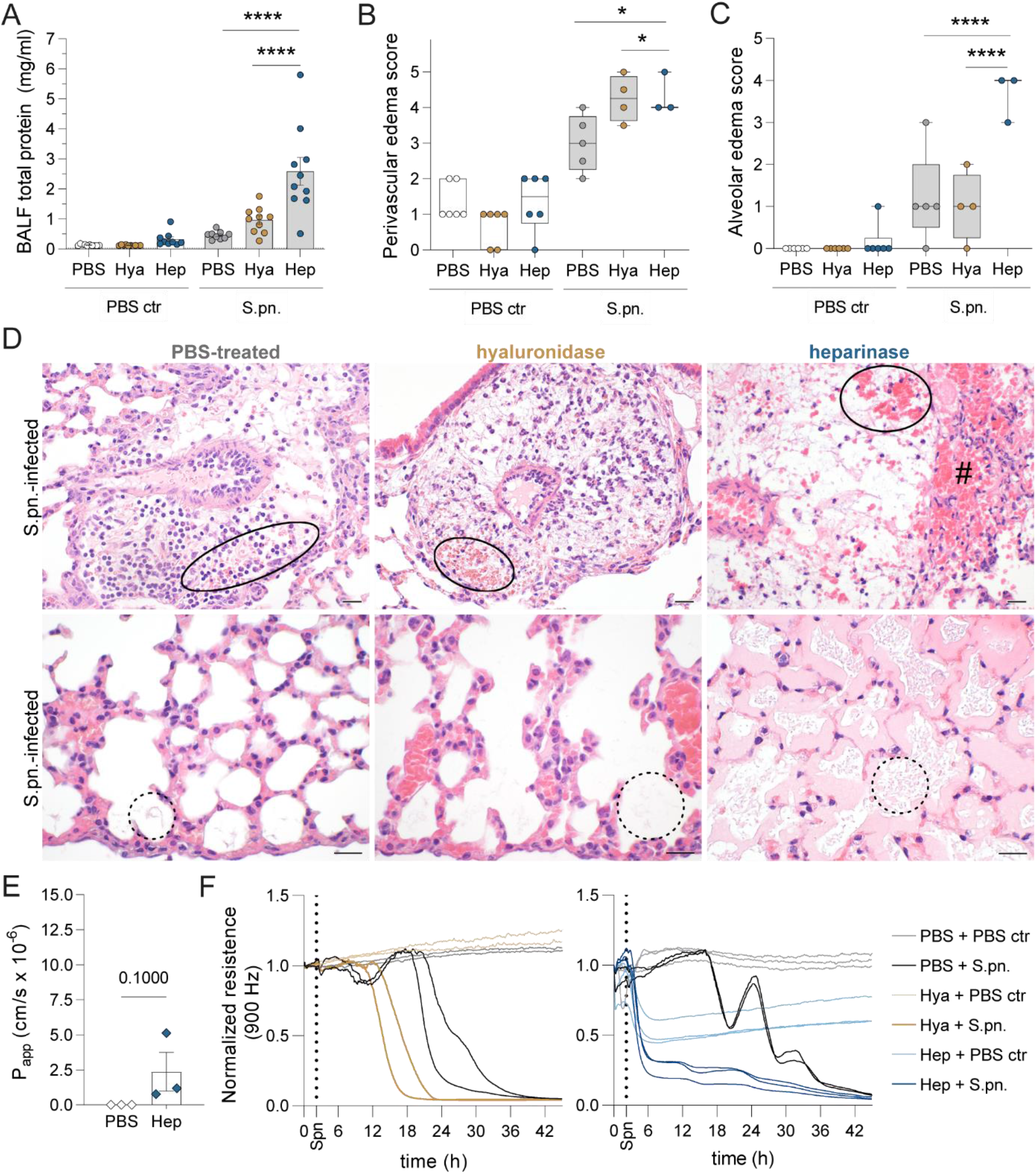
Lung barrier function is impaired by enzymatic glycocalyx modulation during pneumococcal pneumonia. Mice were infected i.n. with *S. pneumoniae* (S.pn.) or PBS (PBS ctr) with addition of hyaluronidase (Hya) or heparinase (Hep) or PBS as control and sacrificed at 48 hpi. (**A**) Protein content in BALF determined by colorimetric assay. Two-way ANOVA, Tukey’s multiple comparisons test; n = 9 – 10. (**B**, **C**) Graph displaying results of histopathological scoring of (**B**) perivascular edema and (**C**) alveolar edema in H&E stained lung sections. Scoring 0 = none, 1 = scattered, 2 = low grade, 3 = medium grade, 4 = severe, 5 = massive. Two-way ANOVA, Tukey’s multiple comparison test. Data are displayed as box plots, middle line displays median, box indicates first and third quartile, and whiskers minimum to maximum; n = 3 – 6. (**D**) Representative H&E staining of murine lungs. Upper panels display representative perivascular edema, ovals highlight perivascular hemorrhage and # highlights alveolar hemorrhage only found in heparinase-treated groups. Lower panels display representative alveolar edema in corresponding groups indicated by dotted circles. Original magnification 400x (upper panel) or 600x (lower panel). Bars equal 20 µm. (**E**) Determination of Alveolus-on-a-Chip barrier integrity upon heparinase treatment by apparent permeability (P_app_) measurement of tracer molecules. Experiment was performed once in triplicate. Mann-Whitney U test. (**F**) TEER measurement by ECIS to assess barrier function of human primary alveolar epithelial cells following hyaluronidase or heparinase or PBS treatment with subsequent S.pn. infection or mock infection (PBS ctr). One representative experiment of n=2-3 shown. * p ≤ 0.05, ** p ≤ 0.01, *** p ≤ 0.001 and **** p ≤ 0.0001.

To determine the direct effects of transnasal fluid inoculation and enzymatic treatments on the alveolar glycocalyx structure, we examined alcian blue-stained tissue samples with transmission electron microscopy at 2 hpi and compared them to samples from untreated WT mice (Fig. E5A, B). Following heparinase treatment, the electron-dense alcian blue layer seemed to be unchanged, while after hyaluronidase treatment we observed areas that were less electron-dense, compared to PBS-treated (Fig. E5A) or non-treated lungs (Fig. E5B). After bacterial infection, the PBS-treated samples showed disorganized filamentous stained material. After infection and heparinase treatment, we observed a denser layer on top of and around the microvilli, suggestive of proteinaceous material coming from extravascular fluid. Interestingly, infected hyaluronidase-treated samples showed less dense but more staining around the microvilli than that observed with the enzymatic treatment alone (Fig. E5A). Thus, we conclude that *in vivo* the applied dose of enzyme treatment does not directly lead to breakdown of the epithelial barrier.

Using human *in vitro* models, we further investigated the role of glycosaminoglycans in regulating lung barrier function. In an alveolus-on-a-chip model (Fig. E6A), we detected minimal tracer translocation to the epithelial channel following apical heparinase treatment, along with signs of epithelial damage whilst the endothelium remained intact (Fig. 6E, E6B). Electric cell-substrate impedance sensing (ECIS) of human primary alveolar epithelial cells (HPAEC) revealed that hyaluronidase treatment alone did not affect epithelial barrier function, whilst heparinase treatment alone did. In combination with bacterial stimulation, pre-treatment with either enzyme resulted in accelerated disruption of the epithelial barrier (Fig. 6F).

## DISCUSSION

In this study, we demonstrate the significant role played by the glycocalyx in limiting the development and severity of pneumococcal disease in mice. Intranasal inoculation with *S. pneumoniae* in the presence of hyaluronidase or heparinase caused more severe disease, resulting in increased pulmonary bacterial burden and inflammatory leukocyte recruitment. Targeting hyaluronan and heparan sulfate resulted in histopathological exacerbations which were manifested in perivascular and intra-alveolar compartments, respectively. Heparinase treatment culminated in increased alveolar permeability *in vivo* and disruption of the epithelial barrier in primary alveolar epithelial cells *in vitro*. Additionally, *Streptococcus pneumoniae* infection induced shedding and desulfation of heparan sulfate in bronchoalveolar fluid.

Hyaluronan and heparan sulfate are the most abundant non-sulfated and sulfated GAGs found in the lungs, respectively, and are involved in a wide spectrum of physiological processes that affect tissue homeostasis (27). Heparan sulfate is a major contributor to the negative charge harbored by the glycocalyx (28), enabling retention of the positively charged chemokines (29) while also playing a role in masking of important leukocyte adhesion molecules such as selectins (30). Hyaluronan can act as a cell adhesion molecule and regulate intercellular and cell-extracellular matrix interactions via its receptor CD44 (31). It is able to bind large volumes of water and has been shown to regulate extravascular lung water volume (32) as well as pulmonary edema during experimental lung injury (33). The precise roles of these glycosaminoglycans in regulating pneumococcal virulence, pulmonary inflammation and epithelial barrier function during bacterial pneumonia, however, remain to be elaborated.

Our findings are in agreement with previous studies investigating hyaluronan in the context of lung edema and streptococcal infections. *S. pneumoniae* has evolved the ability to hydrolyze hyaluronan in order to orchestrate invasion (25), while Group A *Streptococci* can utilize hyaluronic acid capsular polysaccharide to bind epithelial CD44 during colonization of the mouse pharynx (34). Heparan sulfate has previously been shown to interact with *S. pneumoniae* (35) as well as various other respiratory pathogens, including SARS-CoV-2 (36) and *S. aureus* (37). Regardless of which glycosaminoglycan was targeted, the mice in our study exhibited significantly higher bacterial burden in the lungs, coupled to intra-alveolar and luminal growth of pneumococci along with abundant and diffuse distribution of pneumococci within the adventitia of lung vessels.

The majority of histopathological findings such as hemorrhages, neutrophilic infiltrates and edema seen in intra-alveolar and luminal spaces were more prominent in the heparinase-treated group, whilst hyaluronidase treatment could be distinguished by pronounced effects on the lung vasculature. This is in line with the prominent role of intact pulmonary hyaluronan for maintaining vascular homeostasis and preventing leakiness (38). Accordingly, we detected increased splenic bacterial burden, perivascular edema, adventitial neutrophil extravasation and significantly elevated pro-inflammatory cytokine levels in the plasma of hyaluronidase-treated mice.

Heparinase treatment, on the other hand, gave rise to elevated chemokine and cytokine levels in the BAL fluid, including CXCL1 and CXCL5, IL-6 and TNF-α. Considering that endothelial heparan sulfate may bind to and regulate transcytosis of chemokines such as IL-8 (29), it is likely that the intact glycocalyx also binds and stores neutrophil chemoattractants. In addition, it is known that heparan sulfate can act as a ligand for neutrophil L-selectin in order to regulate tissue extravasation (39). Studies have shown that intact, unfractionated heparin binds to and potently inhibits binding of L- and P-selectin to other ligands, while fractionated, low molecular weight heparin molecules lose this ability (40). Accordingly, we observed that treatment with heparinase, which cleaves sulfated saccharides from heparin (41), resulted in the highest number of neutrophils infiltrating into alveolar spaces. Furthermore, heparinase treatment also increased alveolar permeability during pneumococcal pneumonia, a finding we could reproduce *in vitro* in models of epithelial barrier function (alveolus-on-a-chip and ECIS). This is consistent with observations in a murine model of LPS-induced lung injury, in which increased permeability is caused by active shedding of heparan sulfate (9). Thus, targeting heparan sulfate may have a two-pronged effect on neutrophil chemotaxis, by enhancing the availability of chemokines and adhesion molecules, while also increasing epithelial permeability.

The recent demonstrations that alveolar heparan sulfate can be actively shed into the airways during lung inflammation in ARDS patients (9), in an LPS-mediated experimental model of acute lung injury (9) and following influenza infection in mice (42) are in line with our observations. We observed increased GAG shedding upon heparinase treatment, but only in infected mice, indicating that it is an active process driven by pneumococcal pneumonia. In addition, we found that in contrast to control mice, the BAL fluid of infected mice contained predominantly unsulfated heparin disaccharides. While it is possible that pneumococci are directly involved in desulfation of heparin disaccharides through bacterial sulfatases, heparin desulfation was also described in an LPS-mediated acute lung injury model in mice in the absence of live bacterial infection (9), indicating that remodeling of heparan sulfate can be mediated not only by microbes but also by the host. The removal of highly sulfated moieties may further exacerbate infection-induced pulmonary inflammation, as it has been shown that only unfractionated, sulfated heparin is able to block selectin-mediated cell adhesion (43). Furthermore, a recent study demonstrated that highly sulfated motifs of heparan sulfate also inhibit pneumococcal adhesion to host extracellular matrix, resulting in attenuation of corneal infection (44). As some pneumococcal strains may express putative sulfatases in genomic islands (45), it is possible that *S. pneumoniae* sulfatases may contribute to remodeling of heparan sulfate in order to reduce the fraction of sulfated moieties in favor of its virulence. One limitation of our study is the lack of in-depth transcriptional analysis of *S. pneumoniae* during inflammatory conditions, particularly in relation to expression of sulfatases. Unfortunately, the published draft genome sequence of the *S. pneumoniae* serotype 3 NCTC7978 we used in this study (46) lacks the sulfatase genes denoted in the genomic island shown to be present in the *S. pneumoniae* serotype 3 strain (OXC141, ST180).

In conclusion, our study shows the critical importance of pulmonary hyaluronan and heparan sulfate in controlling pneumococcal virulence, disease severity and pulmonary inflammation in mice. Further investigations into the mechanisms of GAG shedding and remodeling, especially in the context of host-pathogen interactions, would provide valuable contributions to infectious disease research and respiratory medicine.

## ACKNOWLEDGMENT

The authors would like to thank Ulrike Behrendt, John Horn and Katja Doerfel for excellent technical assistance.

