## Supplemental Material for "Enzymatic modulation of the pulmonary glycocalyx alters susceptibility to *Streptococcus pneumoniae*"

* Equal contribution

**[#]Corresponding author**

Charitéplatz 1

10117 Berlin

Germany

Tel. + 49 30 450 553347

### EXTENDED MATERIAL AND METHODS

***Bacterial preparation and bacterial growth inhibition assay under enzyme treatment***

*S. pneumoniae* (serotype 3, NCTC 7978; serotype 2, NCTC 7466) was cultured on blood agar plates at 37 °C for 8 h, transferred to liquid medium (Todd Hewitt yeast broth, BD, Heidelberg, Germany, with 10% FCS, Merck, Darmstadt, Germany), stored on ice overnight and grown in a water bath at 37 °C to an $OD_{600nm}$ of 0.35 to 0.40 over 2 to 3 h. For the growth inhibition assay, 60 U heparinase (Heparinase I and III Blend, Merck, Darmstadt, Germany) or 360 U hyaluronidase (Merck, Darmstadt, Germany) were added to 2 mL serotype 3 bacteria in liquid medium and $OD_{600nm}$ was measured in 30 min intervals until stationary phase. For murine infection, serotype 3 bacteria were used. To prepare the infection inoculum, bacteria were washed and adjusted to a concentration of $5 \times 10^8$ colony-forming units (CFUs) per mL in 1×PBS (Thermo Fisher Scientific, Waltham, MA, USA) and mixed 1:2 with 1×PBS, hyaluronidase (90 U in 10 µL 1×PBS), or heparinase (15 Sigma U in 10 µL 1×PBS). The inoculum dose was controlled by CFU counting of serial dilutions.

***Transmission electron microscopy***

For TEM analysis, animals were perfused via the *vena cava* with 0.9% NaCl followed by fixative (1.5% paraformaldehyde (PFA; Electron Microscopy Sciences, Hatfield, United States) and 1.5% glutaraldehyde (GA; Serva, Heidelberg, Germany) in HEPES buffer (Merck Millipore, Darmstadt, Germany), adjusted to pH 7.35). Afterwards, lungs were ventilated through an intratracheally installed venous catheter (22 G), tied and carefully removed, then stored in fixative followed by cutting into small pieces. Randomly chosen pieces were used for further processing. To preserve and visualize the glycocalyx by TEM, we used the cationic agent alcian blue (AB) (1). AB was gently massaged into the lung tissue with a wooden skewer in fixative solution containing 1.5 % PFA with 1.5% GA in 0.15 M HEPES buffer (pH 3) and 0.15% AB 8GX (Sigma Aldrich, St. Louis, Missouri, USA). Finally, the pieces were stored in the fridge in fixative solution with 0.15% AB 8GX for one week to saturate binding sites. Samples were post-fixed using 1% $OsO_4$ (Electron Microscopy Sciences, Hatfield, United States) in 0.1 M cacodylate buffer (Serva, Heidelberg, Germany) at room temperature for 2 h, followed by incubation in half-saturated aqueous uranyl acetate (Serva, Heidelberg, Germany) overnight at 4 °C. After dehydration in a graded acetone series (Roth, Karlsruhe, Germany), samples were transferred to Epon resin (Serva, Heidelberg, Germany). Finally, ultrathin sections of 70 nm were cut using an ultramicrotome (Leica, Wetzlar, Germany) and a diamond knife (Diatome, Nidau, Switzerland), collected on pioloform-coated copper grids (Plano, Wetzlar, Germany) and stained with lead citrate (Merck Millipore, Darmstadt, Germany) according to a published protocol (2). Grids were imaged with a Zeiss Leo 906 electron microscope (Carl Zeiss, Oberkochen, Germany) at 80 kV acceleration voltage equipped with a slow scan 2 K charge-coupled device (CCD) camera (TRS-Tröndle, Moorenweis, Germany) with associated software Image SP (version 1.2.890).

#### *Blood sampling*

Blood was collected during dissection at 48 h from the *vena cava* using a 27 G cannula and 1 mL syringe and transferred into EDTA tubes. Serial dilutions were spread on blood agar plates to determine CFUs. Blood plasma was obtained by centrifugation (2000 × g for 10 min at 4 °C) and stored at -80 °C for further analysis of cytokines and chemokines and for MS to detect shed GAGs.

#### *Bronchoalveolar lavage and organ sampling*

Lungs were lavaged twice through a cannula in the trachea with 0.8 mL PBS containing cOmplete™ Mini protease inhibitor (Merck, Darmstadt, Germany). The bronchoalveolar lavage (BAL) was collected, centrifuged (470 × g for 5 min at 4 °C) and the supernatant BAL fluid (BALF) stored at -80 °C for further analysis. The pellet of BAL cells (BALC) was resuspended in FACS buffer (1×PBS / 0.2% bovine serum albumin, Merck, Darmstadt, Germany) and prepared for FACS analysis. The right lung lobes (superior, middle and inferior lobe) and the spleen were homogenized in 1 mL 1×PBS using a gentleMACS™ Dissociator (Miltenyi Biotec, Bergisch Gladbach, Germany) and serial dilutions spread on blood agar plates to determine CFUs.

#### *Flow cytometry*

BAL cells were blocked against non-specific antibody binding with anti-CD16 / anti-CD32 (BD-Bioscience, Heidelberg, Germany) in FACS buffer for 5 min followed by staining with monoclonal antibodies against CD45, CD11c, Ly6G, SiglecF, (all BD-Bioscience, Heidelberg, Germany), CD11b, F4/80, MHCII (all Thermo Fisher Scientific, Waltham, MA, USA) or Ly6C (BioLegend, San Diego, California, USA) for 20 min at 4 °C in the dark. Cells were washed with 1× PBS, centrifuged (470 × g for 5 min at 4 °C) and fixed overnight in 1% formaldehyde/PBS at 4 °C in the dark. Samples were washed with 1× PBS, centrifuged (470 × g for 5 min, 4 °C) and resuspended in 100 µL FACS buffer before measurements using a FACS Canto II (BD, Heidelberg, Germany). Total cell numbers were calculated using CountBright™ Absolute Counting Beads (Thermo Fisher Scientific, Waltham, MA, USA) according to the manufacturer's instructions. Cells were analyzed using DIVA (BD, Heidelberg, Germany) and FlowJo™ v10.8.1 Software (BD, Heidelberg, Germany) softwares. The gating strategy for the analysis of innate immune cells is shown in Fig. E2.

#### *Colorimetric assays*

For the measurement of total protein, we mixed BALF samples and BSA standards with Protein Assay Reagent S 5000115 and A 5000113 (both Bio-Rad, California, USA), added Protein Assay Reagent B #5000114 (Bio-Rad, California, USA), incubated it for at least 1 h and measured the absorbance at 750 nm using a Multiskan FC Microplate Photometer (Thermo Fisher Scientific, Waltham, MA, USA).

CXCL1 and CXCL5 levels were measured in BALF and plasma using DuoSet ELISA kits (R&D Systems, Inc., Minneapolis, Canada). Other cytokine / chemokine levels were measured in BALF and plasma with LEGENDplex™ Mouse Inflammation Panel (BioLegend, San Diego, California, USA). ELISAs were performed according to the manufacturer's instructions.

To stain GAGs in BALF, the dimethylmethylene blue (DMMB) assay was adapted from published protocols (3) and (4). We used 1,9-dimethylmethylene blue-chloride (16 mg/L, SERVA, Heidelberg, Germany), 3.05 g/L glycine, 1.6 g/L NaCl and acetic acid in distilled water at pH 3.0 for staining and chondroitin sulfate sodium salt (Thermo Fisher Scientific, Waltham, MA, USA) as standard. Absorbance was immediately measured at 525 nm.

***Histopathology, immunohistochemistry***

Dissected lungs were immersion fixed overnight in 4% buffered formaldehyde phosphate solution (neoFroxx, Einhausen, Germany). Sections of 2 µm thickness were cut from paraffin-embedded samples, stained with hematoxylin and eosin (H&E) or prepared for immunohistochemistry (IHC). For *S. pneumoniae* IHC staining, sections were deparaffinized, and endogenous peroxidase was blocked with 10% $H_2O_2$ (pH 7.4) for 15 min. Antigen retrieval was performed by microwaving at 600 W in 750 ml buffered citric acid (pH 6.0) with 1% Triton X 100 (Roth, Germany) for 12 min. Sections were incubated overnight with polyclonal rabbit anti *S. pneumoniae* antibody SPN (diluted 1:500, kind gift from Sven Hammerschmidt, Universität Greifswald, Germany) at 4 °C. Non-specific binding was blocked with 20% goat serum for 30 min. After washing with PBS/Triton buffer, sections were incubated with secondary goat anti-rabbit antibody (diluted 1:200, Vector Laboratories, USA) for 30 min. The signal was enhanced with an ABC-AP kit (Vectastain, USA) and staining was developed with New Fuchsin (Sigma-Aldrich, Germany) for 20 min. Hematoxylin was applied as counterstain. For cellular and bacterial scoring, three 600 × high power fields per compartment (adventitial, intra-alveolar, luminal airway, region with severe pneumonia, region with mild pneumonia, pleural) were analyzed and averaged. Histopathological classification of pneumonia grade, character and localization, based on a grading from 0 (absent) to 4 (severe) (see Table E2), was carried out in a blinded fashion.

***Culture of lung epithelial cells and ECIS***

Human primary alveolar epithelial cells (HPAECs, Cell Biologics Company, Chicago, USA) were grown in Complete Human Epithelial Cell Medium (Cell Biologics Company, Chicago, USA). Passaging of HPAECs was performed by washing with 1xPBS, incubation with TrypLE™ Express (Thermo Fisher Scientific, Waltham, MA, USA) for 7 min at 37 °C for cell detachment, followed by centrifugation (300xg, 5min) and re-seeding at a splitting ratio of 1:3-1:4 in cell culture flasks pre-treated with gelatin-based coating solution (Cell Biologics Company, Chicago, USA). HPAECs between passage 7 and 15 were used for measuring barrier function in ECIS experiments. An ECIS 8W10E+ PET array or a 96W10idf array (ibidi GmbH, Gräfelfing, Germany) was treated with 1× Attachment

Factor Solution (Thermo Fisher Scientific, Waltham, MA, USA) and incubated for at least 30 min at 37 °C. Each well was seeded with 150,000 cells or 50.000 cells in complete medium in the 8W10E+ or 96W10idf array, respectively, and incubated at 37 °C on the ECIS station (ibidi GmbH, Gräfelfing, Germany). Cells were grown on the array until they reached confluence and an electrical resistance of at least 8000 Ohms measured at 900 Hz. Before enzymatic treatment and infection, medium was removed completely and cells were washed two-three times with warm 1xPBS to remove all traces of antibiotics. Then epithelial cell medium without FCS and antibiotics, but with 4 mM of $CaCl_2$ for proper heparinase enzyme activation, was added and cells were incubated for at least 1 h. Next, enzymatic treatment was started. For Hyaluronidase treatment in the 8W10E+ array 25 µl of enzyme stock solution (9000 U/ml, Merck, Darmstadt, Germany) were added to 125 µl medium adding up to a final volume of 150 µl with a concentration of 1500 U/ml. In the 96W10idf array the final volume was 50 µl per well. Heparinase stock solution (1500 Sigma U/ml, Heparinase I and III Blend, Merck, Darmstadt, Germany) and hyaluronidase stock solution were either added directly or pre-diluted in Tris/HCL buffer with 4mM $CaCl_2$ or PBS, respectively, to reach the calculated final concentration (1500 U/ml hyaluronidase; 62.5 Sigma U/ml heparinase) . The controls were treated with either PBS or Tris/HCL buffer. After addition of the enzyme, the array was placed back on the ECIS station and 2 h post-enzyme treatment infection was performed: serotype 2 *S. pneumonia* were diluted in PBS so that cells were infected with 10 µl or 5µl of bacterial suspension at a multiplicity of infection (MOI) 25. Cells were placed back on the ECIS station and measurements resumed for 48 h. Enzyme- and infection-induced disruption of epithelial barrier function was evaluated by normalized values of resistance recorded post stimulation.

#### *Human Alveolus-on-a-Chip*

The Alveolus-on-a-Chip used here is based on the microfluidics organ chip system by Emulate Inc. (Boston, MA, USA). The protocol was adapted from (5). In brief, after activation of the chips with ER1/ER2 components as per manufacturer's instructions, the chips were coated with a mixture of collagen IV (200 µg/ml, Sigma, C5533) and laminin (15 µg/ml, Sigma, L6274) overnight at 4 °C. On day 0, pre-passaged human primary alveolar epithelial cells (HPAECs, Cell Biologics, H-6053, Lot # F101517Y72) and human pulmonary microvascular endothelial cells (HPMVECs, Promocell, C-12281, Lot # 474Z031.1 and 463Z013) were sequentially seeded on the chip with at least 2 h between seeding the different cell types. HPAECs were seeded in complete human epithelial cell medium (Cell Biologics, H6621) and HPMVECs in complete EGM-MV2 (Promocell, C-22121) with seeding densities of $1.6 \times 10^6$ cells/ml and $8 \times 10^6$ cells/ml, respectively. After cell attachment, the chips were washed by gravity flow and kept under static conditions at 37 °C, 5% $CO_2$. The next day (day 1), the chips were connected to the Pods (Emulate Inc.) and the Pods placed into the Zoe instrument (Emulate Inc.). The instrument was set to perfuse the top and bottom channel of the chips at 60 µl/h. On day 4, dexamethasone (1µM, Sigma, D4902) was added to the epithelial cell medium of the top channel to strengthen tight junctions. Two days later (day 6), air-liquid interface culture was induced by flushing the apical epithelial channel at

1,000 µl/h for 5 min. Subsequently, all cells on the chip were only fed from the endothelial channel with EGM-MV2. On day 8, FCS in the EGM-MV2 medium was reduced from 5% to 0.5%. On day 11, cyclic stretch was applied to the chip membrane at 5% strain and 0.25 Hz to recapitulate a physiological breathing function. Chips with an intact barrier (no leakage in the apical channel) were used on day 15 for enzymatic heparinase treatment (final concentration of 750 Sigma U/ml in PBS, Heparinase I and III Blend, Merck, Darmstadt, Germany). The control was treated with PBS only. The next day, barrier function was analyzed by applying tracer (Cascade Blue™ hydrazide, Trisodium Salt, Invitrogen, C687) from the bottom endothelial channel for 2 h (120 µl/h) and collecting outflows from the apical epithelial channel as well as the bottom channel. Fluorescence was measured with a Spectra Max M2 plate reader (Molecular Devices) and apparent permeability ($P_{app}$) was calculated as described previously (6).

For immunofluorescence staining, chips were either cut in half or stained as a whole after fixation with 3.7% paraformaldehyde for 20min at room temperature. After fixation, chips were washed twice with 1x PBS, permeabilized for 30 min at room temperature (1% Saponin/PBS), blocked for 2 h at room temperature (10% donkey serum and 1% BSA/PBS) and washed 3x with 200 µl 1xPBS. Primary antibodies were diluted in 1% BSA/PBS and incubated at 4 °C overnight. The bottom channel was perfused with mouse anti VE-Cadherin antibody (Santa Cruz, sc9989, 1:100) to stain endothelial junctions, while the top channel was exposed to mouse anti HTII-280 antibody (Terrace, TB-27AHT2-280, 1:200) marking alveolar type 2 cells. After washing 3x with 200 µl 1xPBS, chips were incubated with secondary antibodies (donkey anti-mouse IgG Alexa Fluor™ 488, Thermo Fisher, A-21202, 1:1000) for 2 h at room temperature. Washing was followed by incubation with DAPI (Abcam, ab228549) for 15 min at room temperature and final washing (3x with 200 µl 1xPBS). Stained chips were analyzed with a LSM980 confocal microscope (ZEISS).

***Disaccharide analysis***

BALF and plasma samples were dried completely and resuspended in 50 mM Tris/ 10 mM CaCl2, pH 7.6. Pronase (Sigma-Aldrich Chemie GmbH, Taufkirchen, Germany) was added up to a concentration of 10 mg/ml and samples were incubated overnight at 37 °C. The following day, pronase was inactivated by heating samples at 95 °C for 5 min. After cooling down, 250 U of benzonase (Sigma-Aldrich Chemie GmbH, Taufkirchen, Germany) in 2 mM MgCl2 (VWR International GmbH, Darmstadt, Germany) was added and samples incubated for 4 h at 37 °C. Afterwards, GAGs were enriched using manual anion exchange chromatography with Q-sepharose beads (Cytiva, Marlborough, Massachusetts, USA). Self-packed columns were equilibrated with 20 mM NaOAc/100 mM NaCl (Sigma-Aldrich Chemie GmbH, Taufkirchen, Germany and VWR International GmbH, Darmstadt, Germany), pH 5.0, before acidified samples were applied. After thoroughly washing with equilibration buffer, the elution was carried out using 20 mM NaOAc/ 1 M NaCl, pH 5.0. After complete elution, samples were desalted using cold, NaOAc -saturated ethanol precipitation. The samples were stored in ethanol (Roth, Karlsruhe, Germany) and incubated overnight at -20 °C followed by centrifugation (21,000 × g for 20 min at 4 °C). The supernatant was removed and pellets were dried completely. Pelleted GAGs were resuspended in 20 mM Tris/5 mM $CaCl_2$ (both Sigma-Aldrich Chemie GmbH, Taufkirchen, Germany)/200 mM NaCl, pH 7.0, and incubated at 30 °C. A heparinase cocktail (10 mU of heparinase 1, 2 and 3, produced in-house) was added for heparin disaccharide analysis. For chondroitinsulfate depolymerization, 10 mU of chondroitinase ABC (Sigma-Aldrich Chemie GmbH, Taufkirchen, Germany) was added in 150 mM NaCl, 10 mM sodium phosphate pH 7.4 (Sigma-Aldrich Chemie GmbH, Taufkirchen, Germany). The total volume of each reaction was 40 µl. After overnight incubation at 37 °C, enzymes were inactivated by heat and samples centrifuged at 21,000 × g for 10 min at 4 °C. The supernatants were transferred and freeze dried, then 20 µl of a 0.1 M 2-aminoacridone-HCl (Sigma Aldrich, St. Louis, Missouri, USA) solution in acetic acid/DMSO (3:17 vol/vol; Sigma-Aldrich Chemie GmbH, Taufkirchen, Germany and VWR International GmbH, Darmstadt, Germany) was added to the pellets. Dissolved samples were incubated for 15 min in the dark at room temperature. After incubation, an equal volume of 1 M sodium-cyanoborohydride (TCI EUROPE N.V., Zwijndrecht, Belgium) was added and samples incubated for 4 h at 45 °C. After incubation, the samples were centrifuged at 21,000 × g at 4 °C for 15 min. The supernatants were collected and freeze dried. Pure acetone (Sigma-Aldrich Chemie GmbH, Taufkirchen, Germany) was added up to a total volume of 500 µl to lyophilized samples. After centrifugation at 21,000 × g for 15 min at 4 °C, the supernatants were removed and pellets were washed one more time with acetone. Precipitated samples were dried completely, dissolved in 3% acetonitrile and then loaded onto an Aquity UPLC BEH C18 column (2.1 × 150 mm, 1.7 µm; Waters Corporation, Milford, Massachusetts, USA) which was installed in a Knauer Azura UHPLC system (KNAUER Wissenschaftliche Geräte GmbH, Berlin, Germany). The mobile phase consisted of 150 mM NH4OAc pH 5.6 (A) and the eluent was 100% acetonitrile (B). The starting conditions were 97% A/3% B. A gradient from 3% B to 13% B within 20 min was applied. The detection was performed at an excitation wavelength of 425 nm and an emission wavelength of 520 nm.

### EXTENDED REFERENCES

1. Behnke O, Zelander T. Preservation of intercellular substances by the cationic dye alcian blue in preparative procedures for electron microscopy. J Ultrastruct Res 1970;31:424–8.

2. REYNOLDS ES. The use of lead citrate at high pH as an electron-opaque stain in electron microscopy. J Cell Biol 1963;17:208–12.

3. Zheng CH, Levenston ME. Fact versus artifact: avoiding erroneous estimates of sulfated glycosaminoglycan content using the dimethylmethylene blue colorimetric assay for tissue-engineered constructs. Eur Cell Mater 2015;29:224-36; discussion 236.

4. Sun X, Li L, Overdier KH, Ammons LA, Douglas IS, Burlew CC, Zhang F, Schmidt EP, Chi L, Linhardt RJ. Analysis of Total Human Urinary Glycosaminoglycan Disaccharides by Liquid Chromatography-Tandem Mass Spectrometry. Anal Chem 2015;87:6220–7.

5. Bai H, Si L, Jiang A, Belgur C, Zhai Y, Plebani R, Oh CY, Rodas M, Patil A, Nurani A, Gilpin SE, Powers RK, Goyal G, Prantil-Baun R, Ingber DE. Mechanical control of innate immune responses against viral infection revealed in a human lung alveolus chip. Nat Commun 2022;13.

6. Si L, Bai H, Rodas M, Cao W, Oh CY, Jiang A, Moller R, Hoagland D, Oishi K, Horiuchi S, Uhl S, Blanco-Melo D, Jordan T, Nilsson-Payant BE, Golynker I, Frere J, Logue J, Haupt R, McGrath M, Weston S, Zhang T, Plebani R, Soong M, Nurani A, Kim SM, Zhu DY, Benam KH, Goyal G, Gilpin SE, Prantil-Baun R, Gygi SP, Powers RK, Carlson KE, Frieman M, tenOever BR, Ingber DE. A human-airway-on-a-chip for the rapid identification of candidate antiviral therapeutics and prophylactics. Nat Biomed Eng 2021;5:815–29.

### SUPPLEMENTAL TABLES

**Suppl. Table E1: Murine clinical disease scoring**

| SCORE | ACTIVITY | BODY CARE | BREATHING |
|---|---|---|---|
| 0 | Curious, awake | Shiny fur | Steady |
| 1 | Less active | Disheveled fur or tearing eyes | Hypo- or hyper-ventilation |
| 2 | Less reactive to stimulation or isolated | Disheveled fur and tearing eyes | Labored |

**Suppl. Table E2: Scoring scheme for grade of pneumonia**

| SCORE | GRADE OF PULMONARY INFLAMMATION |
|---|---|
| 0 | Absent |
| 1 | Minimal |
| 2 | Low-grade |
| 3 | Moderate |
| 4 | Severe |

### SUPPLEMENTAL FIGURES

#### Suppl. Figure E1

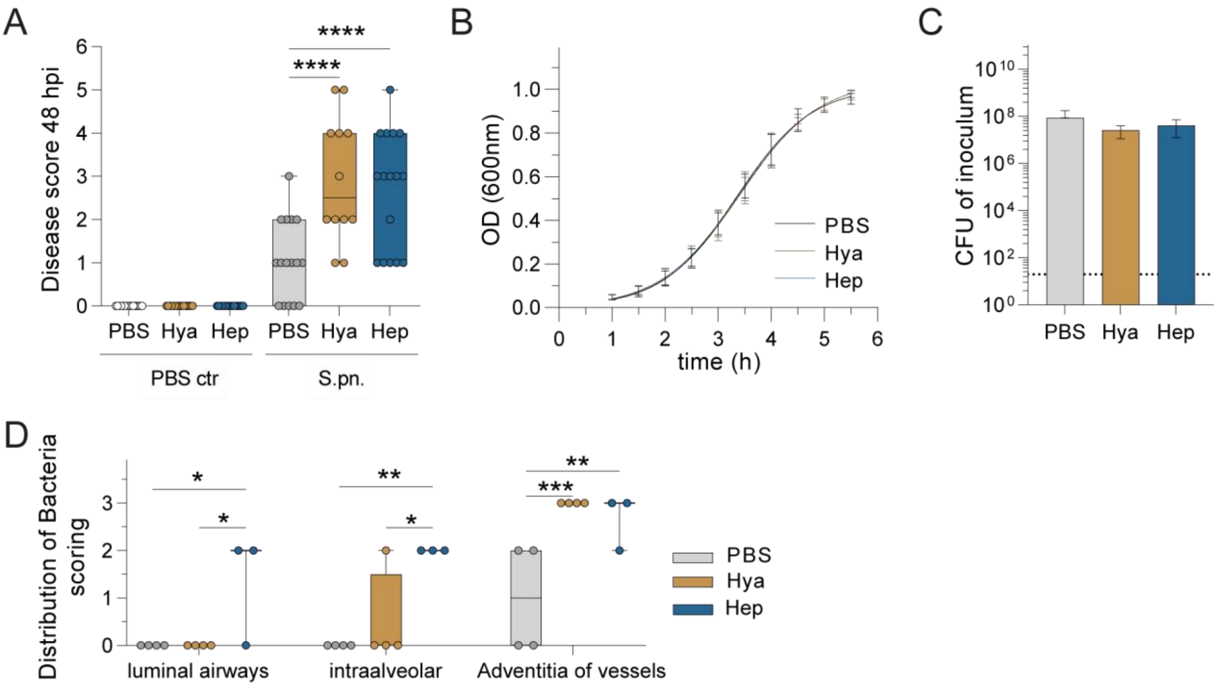


**Figure E1. Murine disease score, *in vitro* bacterial growth curves and bacterial distribution *in situ*.** (**A, D**) Mice were infected i.n. with *S. pneumoniae* with addition of hyaluronidase (Hya) or heparinase (Hep) or PBS as control. Behavioral disease scoring of surviving animals at 48 hpi; n = 10 – 16. (**B**) *In vitro* growth curves and (**C**) CFU from inoculum of *S. pneumoniae* in absence and presence of enzymes; n = 3 for both. LOD, 20 CFU (**D**) Distribution of bacteria. S.pn. staining on lung sections was scored as 0 (absent), 1 (focal), 2 (multifocal) or 3 (diffuse); n = 3 – 4. (**A**, **D**) Two-way ANOVA, Tukey's multiple comparisons test; middle line in box plot displays the median, the box indicates the first and third quartile, whiskers indicate the minimum to maximum. * $p \leq 0.05$, ** $p \leq 0.01$, *** $p \leq 0.001$ and **** $p \leq 0.0001$.

**Suppl. Figure E2**

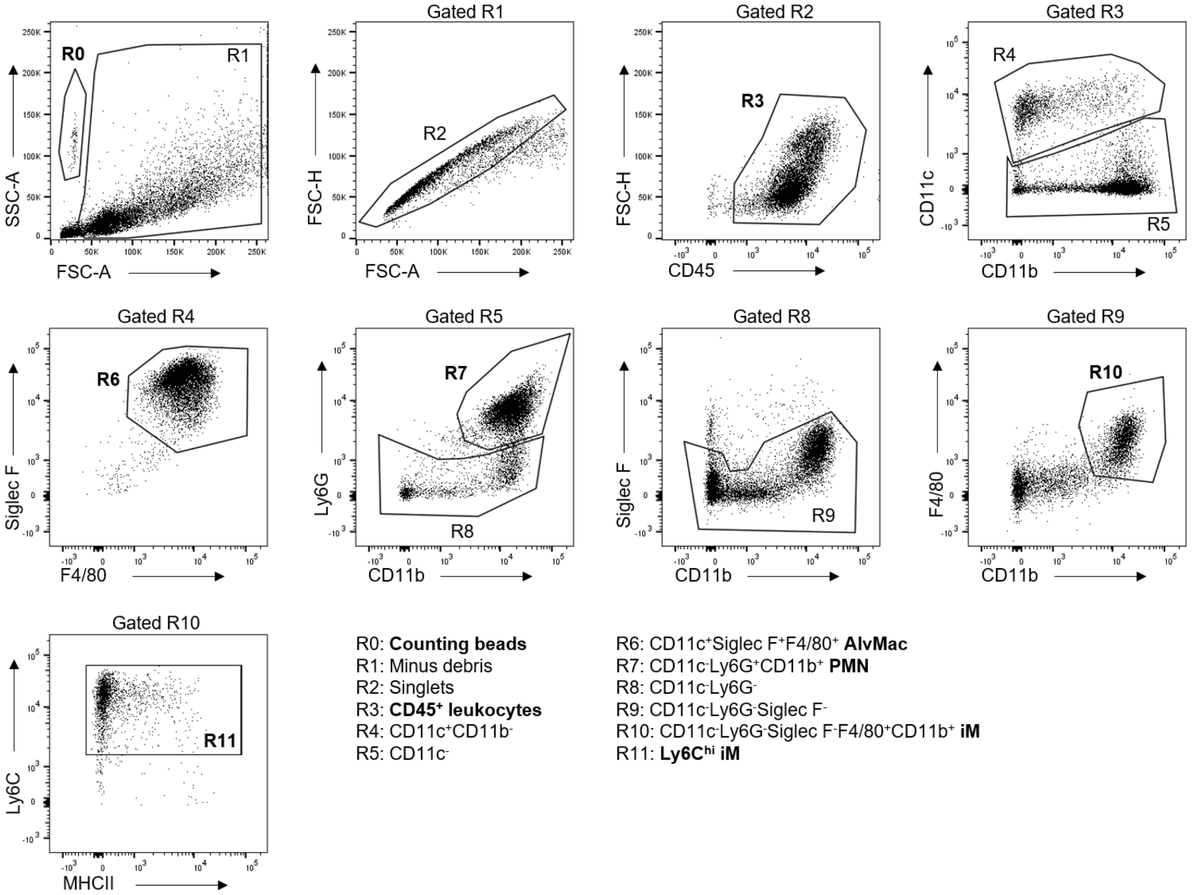


**Figure E2. Flow cytometry gating strategy for identification of innate immune cell populations in BAL.** Forward scatter (FSC) and side scatter (SSC) were utilized to define regions (R); R0 for counting beads and R1 for debris exclusion. Populations were gated on R1, R2 to define singlet population by FSC-area (A) versus FSC-height (H). Gate R2 was used to define R3 as CD45$^{+}$ leukocytes. Gated leukocytes (R3) were used to identify CD11c and CD11b populations, which in turn were used to discriminate gate R4 (CD11c$^{+}$CD11b$^{-}$) and R5 (CD11c$^{-}$). R4 gated leukocytes were further discriminated in R6 by SiglecF$^{+}$F4/80$^{+}$ gating of alveolar macrophages (AlvMac). R5 gated leukocytes were further discriminated in R7 by Ly6G$^{+}$CD11b$^{+}$ gating to identify neutrophils (PMN) and in R8 by CD11c$^{-}$Ly6G$^{-}$ leukocytes. R8 gated leukocytes were used to identify CD11c$^{-}$Ly6G$^{-}$SiglecF$^{-}$ leukocytes (R9). Gated on R9, we identified CD11c$^{-}$Ly6G$^{-}$SiglecF$^{-}$F4/80$^{+}$CD11b$^{+}$ leukocytes (R10), namely inflammatory macrophages (iM) by F4/80 versus CD11b gating. iMs in R10 could be further used to identify Ly6C$^{hi}$ monocyte-derived inflammatory macrophages (Ly6C$^{hi}$ iM; R11).

**Suppl. Figure E3**

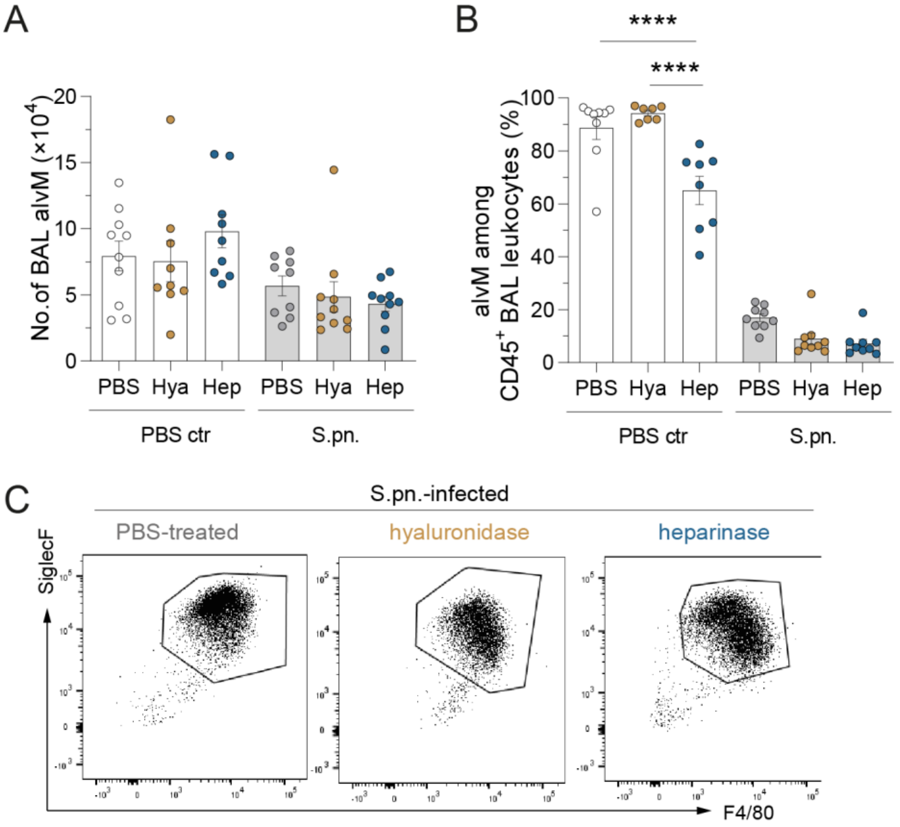


**Figure E3. Alveolar macrophage numbers and proportions in BAL.** Mice were infected i.n. with *S. pneumoniae* (S.pn.) or PBS (mock-infected) with addition of hyaluronidase (Hya) or heparinase (Hep) or PBS as control and sacrificed at 48 hpi. Flow cytometry-based detection and quantification of alveolar macrophages. (**A**) Total numbers of alveolar macrophages in BAL were determined by use of counting beads while (**B**) cell frequencies were determined as a percentage of $CD45^+$ leukocytes. (**C**) Dot plots representing cellular frequencies amongst $CD45^+$ leukocytes. (**A, B**) Two-way ANOVA, Tukey's multiple comparisons test; n = 7 – 10. **** $p \leq 0.0001$

**Suppl. Figure E4**

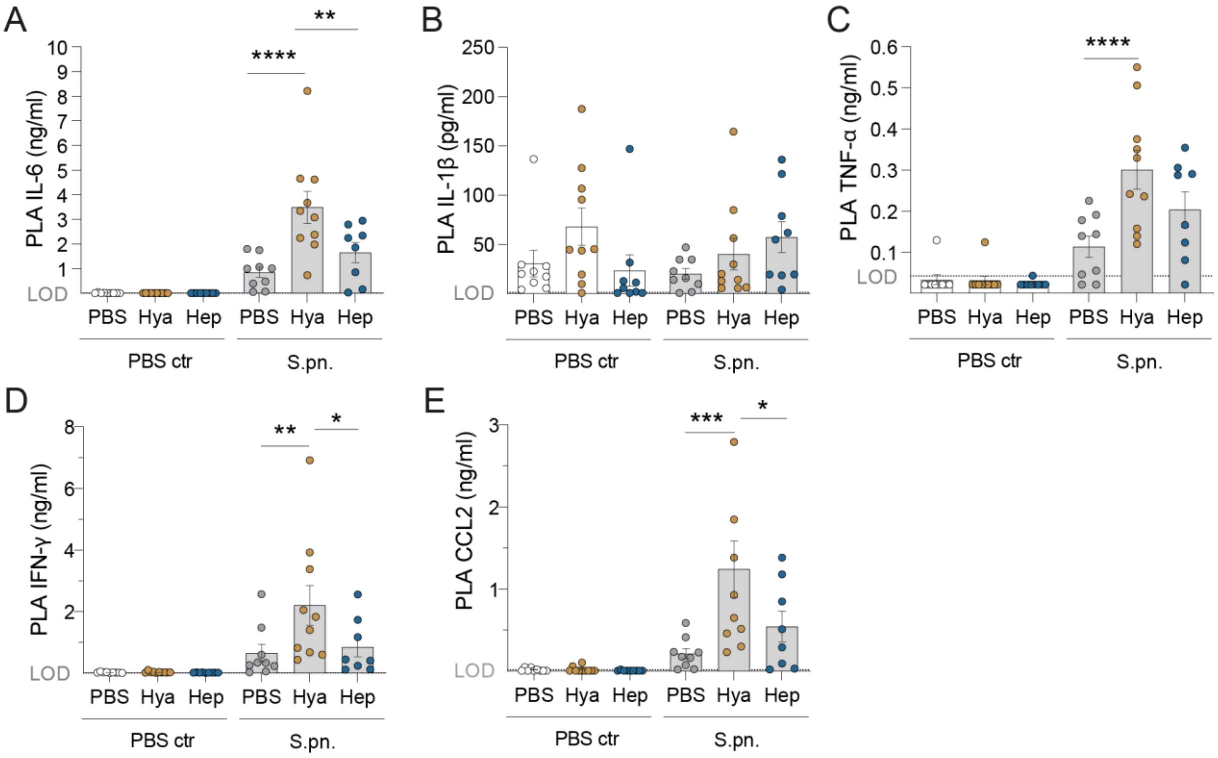


**Figure E4. Pro-inflammatory cytokines and chemokine CCL2 in blood plasma.** Mice were infected i.n. with *S. pneumoniae* or PBS (mock-infected) with addition of hyaluronidase (Hya) or heparinase (Hep) or PBS as control and sacrificed at 48 hpi. The concentrations of pro-inflammatory cytokines and chemokine CCL2 in plasma were measured by ELISA: (**A**) IL-6 (LOD 2.68 pg/ml), (**B**) IL-1β (LOD 1.46 pg/ml), (**C**) TNF-α (LOD 43 pg/ml), (**D**) IFN-γ (LOD 7.7 pg/ml), (**E**) CCL2 (LOD 16.51 pg/ml). One plasma sample from heparinase-treated animals was removed from analysis, as it exceeded values by >10-fold and identified as outlier across all cytokine measurements by ROUT method. Two-way ANOVA and Tukey's multiple comparison test. n = 9 – 10. * $p \leq 0.05$, ** $p \leq 0.01$, *** $p \leq 0.001$ and **** $p \leq 0.0001$.

### Suppl. Figure E5

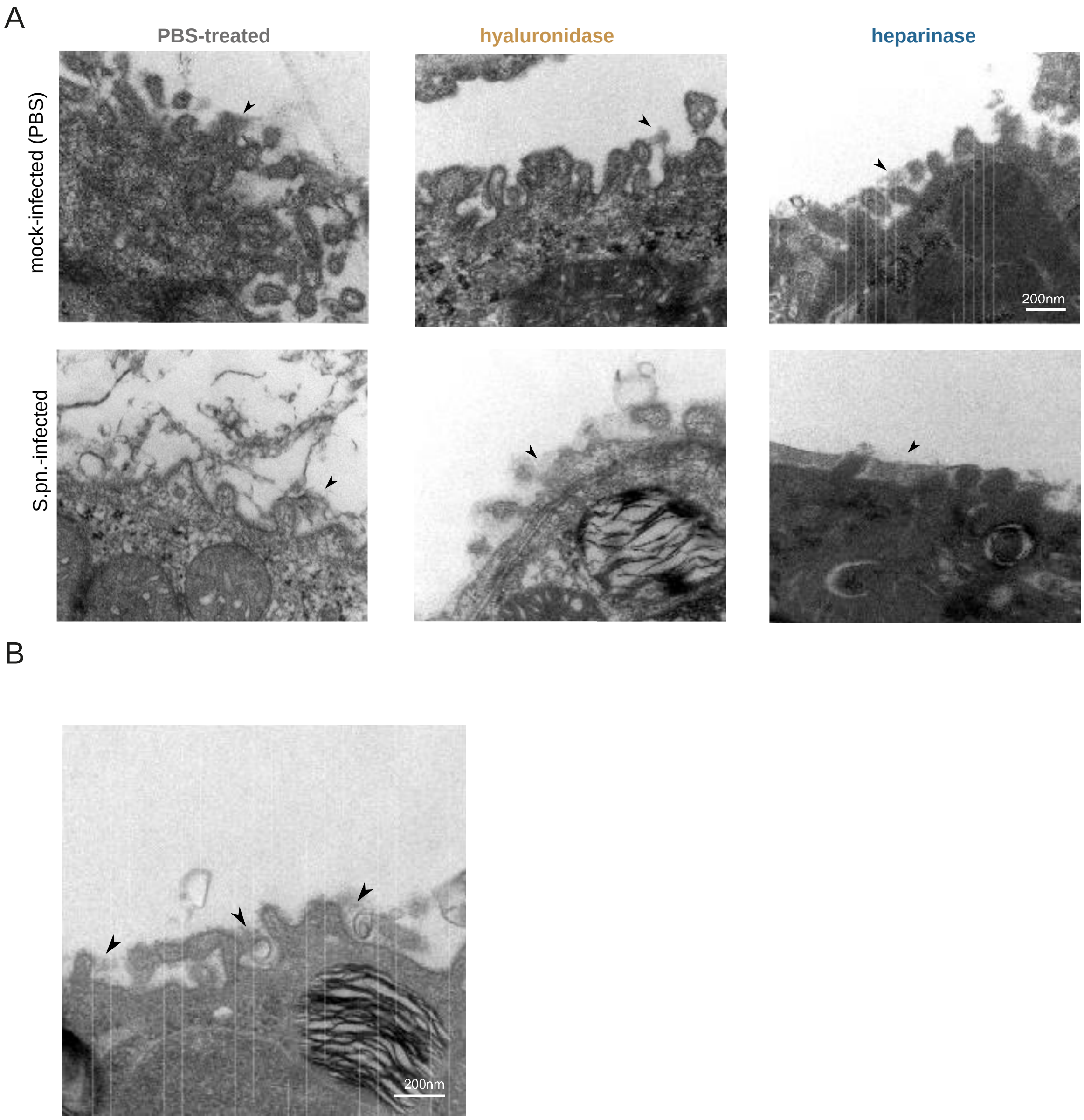


**Figure E5. Representative transmission electron micrographs of alcian blue-stained glycocalyx on alveolar epithelial type 2 cells (AEC2).** (**A**) PBS (left), hyaluronidase (middle) or heparinase (right) treated lungs from mock-infected (top panel) and S.pn.-infected (bottom panel) mice. (**B**) Naïve (non-treated) lungs stained with alcian blue. Arrows indicate the electron-dense alcian blue-stained alveolar epithelial glycocalyx on top and around microvilli on AEC2. Scale bars: 200 nm.

**Suppl. Figure E6**

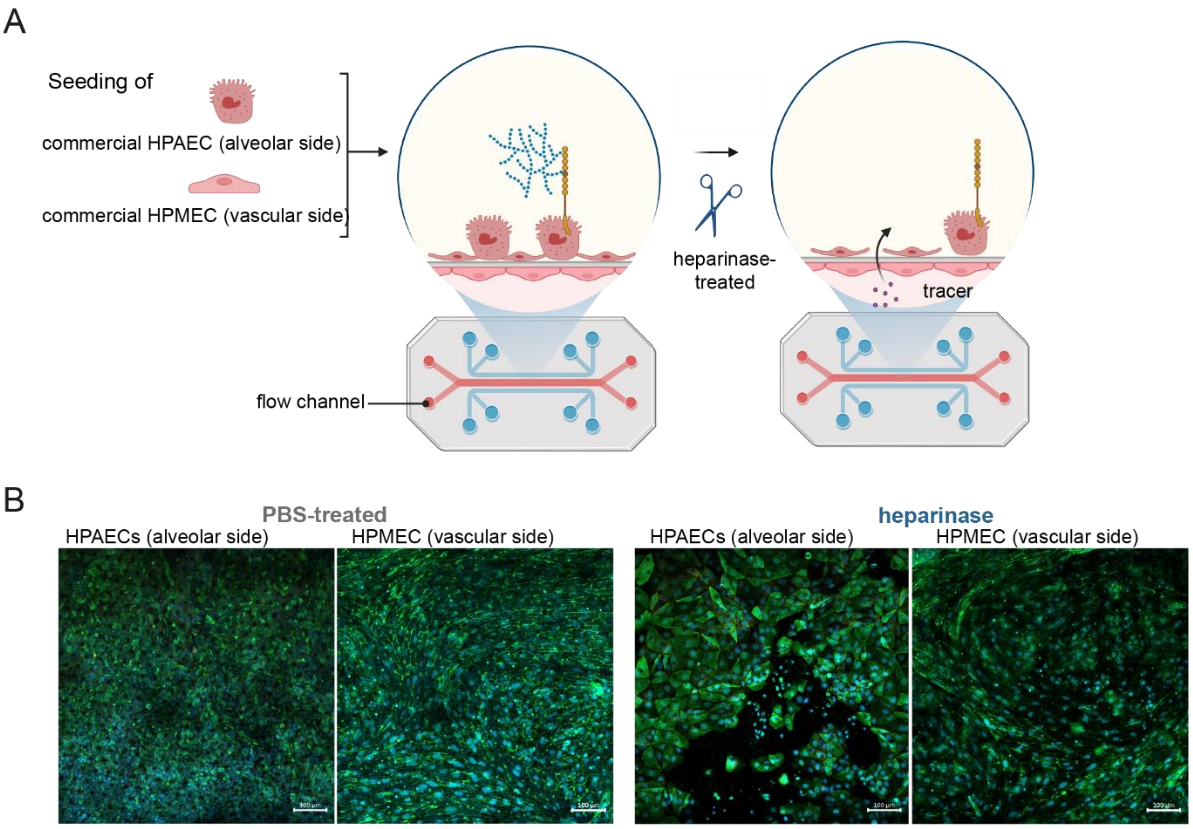


**Figure E6. Heparinase treatment enhances vascular-alveolar permeability in human alveolus-on-a-chip models.** (**A**) Scheme depicting human alveolus-on-a-chip layout and experimental treatment. (**B**) Immunofluorescent staining of human primary alveolar type 2 epithelial cells (HPAECs, HTII-280, green) on the alveolar side and human pulmonary microvascular endothelial cells (HPMECs, VE-Cadherin, green) on the vascular side. Nuclei are stained with DAPI (blue). Scale bars: 100 µm.
